# Short-stranded zein fibers for muscle tissue engineering in alginate-based hydrogels

**DOI:** 10.1101/2023.10.18.562894

**Authors:** Lea Melzener, Sergio Spaans, Nicolas Hauck, André J. G. Pötgens, Joshua E. Flack, Mark J. Post, Arın Doğan

## Abstract

Cultivated meat is a nascent technology that aims to produce an environmentally and animal-friendly alternative to conventional meat. Producing skeletal muscle tissue in an animal-free system allowing for high levels of myofusion and maturation is important for the nutritional and sensorial value of cultivated meat. Alginate is an attractive biomaterial to support muscle formation as it is food-safe, sustainable, cheap, and can be cross-linked using non-toxic methods. Although alginate can be functionalized to promote cell attachment, limitations in its mechanical properties, including form, viscosity and stress relaxation, hinder cellular capacity for myogenic differentiation and maturation in alginate-based hydrogels. Here, we show that the addition of electrospun short-stranded zein fibers increased hydrogel degradation, resulting in faster compaction, improved cell-gel interaction and enhanced alignment of bovine muscle precursor cells. We conclude that fiber-hydrogel composites are a promising approach to support optimal formation of 3D constructs, by improving tissue stability and thus prolonging culture duration. Together, this improves muscle-related protein content by facilitating myogenic differentiation and priming muscle organoids for maturation.

## Introduction

Cultivated meat (also known as ‘cultured meat’) is an evolving field that aims to produce meat products without the adverse environmental and animal welfare implications inherent to conventional meat production^1–3^, by leveraging the *in vitro* proliferation and differentiation of stem cells. Although substantial progress has been made in recent years, including the development of animal component-free processes^4–8^, several major challenges are still being addressed to facilitate mass commercialization of these technologies^3^.

A central premise of muscle tissue engineering for cultivated meat is the emulation of sensory and nutritional properties of skeletal muscle, and is contingent on the myogenic differentiation of muscle precursors (such as myoblasts) to form myotubes in three-dimensional tissue constructs. Subsequent maturation, marked by increased biosynthesis of muscle-related structural proteins, might help mimic the characteristic taste, texture, mouthfeel and nutritional value of conventional meat^9,10^. Maturation of a bioartificial muscle (BAM) necessitates supportive structures that function to promote tissue formation, providing mechanical stability and cell-attachment sites that prolong the survival of myoblasts and myotubes, thereby ensuring extended organoid survival^11^.

Traditionally, animal-derived proteinaceous hydrogel systems, such as collagen and fibrin, have been extensively used for muscle tissue engineering, and were investigated for their applicability for development of potential treatments and drug screening methodologies^12^. As extracellular matrix (ECM) proteins, these hydrogels present cell attachment sites and fibrous constituents, aiding cells in adherence, spreading and migration. Such systems rely on compaction to enable cell-cell contact (which is imperative for myogenic fusion into myotubes) and depend on anchorage to induce tension, which results in cell alignment, priming the construct for differentiation. Development of a uniform tissue in which cells are aligned by tension and guided by fibrous structures, thereby promoting sustained myotube attachment, is an indispensable aspect of tissue maturation^13^. However, these systems are animal-based, and hence cannot be applicable for animal component-free product development. While diverse options for plant-based polysaccharides and decellularized materials have been explored to support BAM formation ^5,14^, their reduced degradability, inadequacy in creating tension, absence of mechanical support and lack of attachment cues for myocytes hamper BAM viability.

Alginate is a commonly employed polysaccharide for muscle tissue engineering, because its tunable mechanical and physical properties are well tailored to cellular requirements, alongside well-established options for peptide functionalization^15^. The presence of calcium (an ionic cross-linker) significantly influences the tissue formation and gel contraction behavior of alginate hydrogels. The dynamic nature of this cross-linking facilitates tunable mechanical properties and degradation of the tissue, adapting to the evolving requirements of the satellite cells, as they spread, differentiate and fuse into myotubes. Nevertheless, alginate-based systems lack fibrous components akin to those available in natural ECM and collagen-based hydrogels, which benefit tissue formation and stability^16,17^. Various studies have attempted to address these limitations, often through the integration of a secondary component. Blending fibrous components has been shown to improve mechanical properties of hydrogels, resulting in hybrid systems termed fiber-hydrogel composites (FHCs)^18^.

Electrospinning provides an attractive method for producing such fibers for hydrogel integration. Electrospun fibers are easy to fabricate and the method is amenable to upscaling, while their fibrous structure can mimic natural ECM constituents, which, in combination with the aforementioned properties of the primary biomaterial, can assist tissue formation and maturation. Prior research has demonstrated convincing proof of principle in animal-based and non-edible systems, for example employing murine C2C12 cells encapsulated within gelatin-based hydrogels, with secondary fibrous compounds fabricated from polycaprolactone (PCL)^19^. Translating these advantages to a cultivated meat system, however, necessitates an animal-free and food-compatible method. To this end, we utilized zein, a heterogeneous inherently hydrophobic protein mixture isolated from maize (*Zea mays*)^20^,. Alpha-zein (a 19 kDa glutamine-rich protein which is the predominant component of zein) has been established as cytocompatible, and is known to be amenable to electrospinning^21, 22^.

In this study, we aimed to enhance the mechanical and physical attributes of an alginate hydrogel to improve cellular adhesion and compaction of 3D skeletal muscle tissues. We achieved this by incorporating electrospun zein fibers, leading to improved cell alignment and scaffold stability to support myogenic differentiation and maturation.

## Results

### Electrospinning of zein fiber solutions (Table 1, Fig. 1, Fig. 2)

In order to produce electrospun zein fibers, which can be processed into short-stranded fibers (SSFs) and incorporated into RGD-functionalized alginate hydrogels (Fig. 1) to improve tissue formation and cell alignment, we first optimized the electrospinning conditions. We characterized and compared 5 different solutions of 20, 25, 28, 30 and 35 wt% zein dissolved in a 1:1 mixture of acetic acid and ethanol (Table 1). We measured the physical properties of these solutions, including conductivity, surface tension and viscosity, which can affect the electrospinning process and fiber morphology.

**Figure 1:**
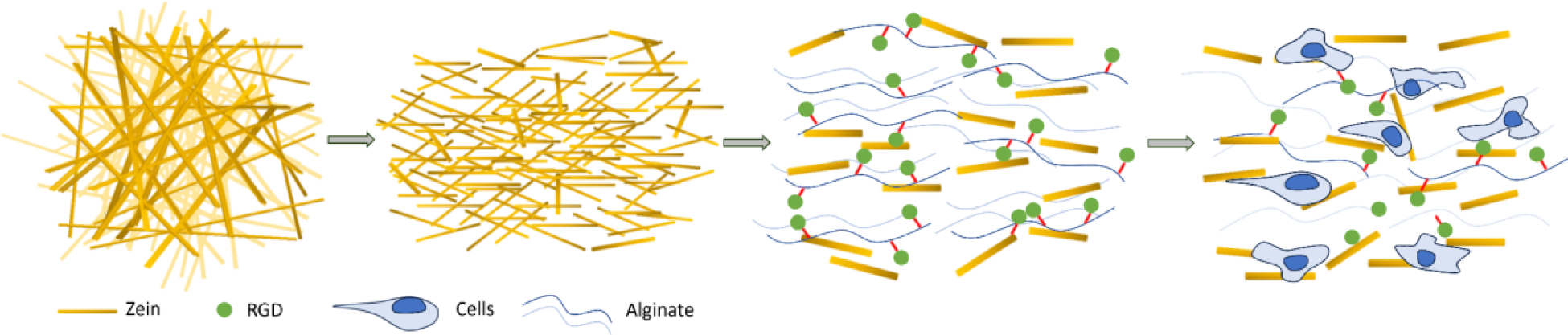
Schematic overview of fabrication and incorporation of short stranded zein fibers (SSFs) into RGD-alginate BAMs.

**Table 1:** Characterization of zein solutions for electrospinning.

| Condition | 20 wt% | 25 wt% | 28 wt% | 30 wt% | 35 wt% |
| --- | --- | --- | --- | --- | --- |
| **Zein concentration [mg ml<sup>-1</sup>]** | 200 | 250 | 280 | 300 | 350 |
| **Viscosity [mPa.s]** | 81 | 208 | 359 | 544 | 1708 |
| **Conductivity [μS cm<sup>-1</sup>]** | 281.2 | 258.2 | 240.0 | 226.8 | 195.6 |
| **Taylor cone formation** | yes | yes | yes | yes | no |
| **Morphology** | bead formation | mostly fibers w/ some bead formation | homogeneous fibers | heterogeneous melted fibers | heterogeneous thick fibers |
| **Fiber diameter [μm]** | 0.130 | 0.365 | 0.664 | 0.980 | 1.815 |
| **Fiber size distribution [μm]** | $\pm 0.029$ | $\pm 0.136$ | $\pm 0.202$ | $\pm 0.639$ | $\pm 0.889$ |

The increased conductivity and lower viscosity of the different solutions with lower zein concentrations correlated with the spinning results that led to thin fiber and bead formation as opposed to the homogeneous fibers we aimed for to produce SSFs (Fig. 2, Table 1). The opposite trend was seen as the zein concentration was increased to 28, 30 and 35 wt%, which resulted in thicker fibers. However, at concentrations higher than 30 wt%, the Taylor cone did not form during the electrospinning process, which led to inhomogeneous fiber morphologies (Fig. 2, Table 1). Since 28 wt% zein proved most amenable to a robust electrospinning process, and yielded the most promising fibrous zein sheets with homogeneous fiber size distribution, we proceeded with this solution for subsequent experiments.

**Figure 2:**
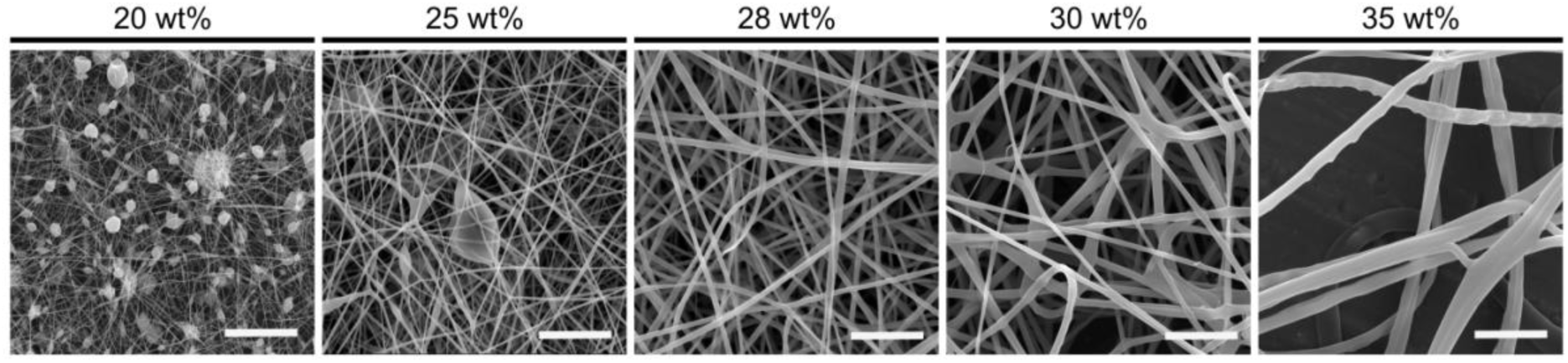
Morphology of electrospun zein fibers. Scanning electron microscopy (SEM) images showing morphology of electrospun fibers using 20, 25, 28, 30 and 35 wt% zein solution. Scale bars, 10 µm.

### Zein fibers can be processed into cytocompatible short stranded fibers (Fig. 3)

After manufacturing electrospun zein fibers, we investigated the cytocompatibility of the fibrous zein sheets in a serum-free cell culture system, using primary bovine satellite cells (SCs). Promisingly, we observed that cells were able to attach and spread on the zein structures within 24 hours without additional peptide functionalization (Figs. 3a, b). This observation suggests that zein protein as a secondary compound adds additional attachment sites to the 3D alginate construct, and thus could improve cell-gel interaction. In order to form FHCs in which the dynamicity of the hydrogel is maintained, further processing of the zein sheets into SSFs was required. We first tested various electrospinning parameters to optimize the alignment of fiber scaffolds, and found that increasing the rotation rate of the collector from 200 to 2500 rpm substantially improved fiber alignment. Both the non-aligned and aligned fibers were processed into SSFs using ultrasonication. We observed that processing of aligned zein scaffolds resulted in reduced clustering of SSFs compared to the non-aligned fibers (Figs. 3c, d).

**Figure 3:**
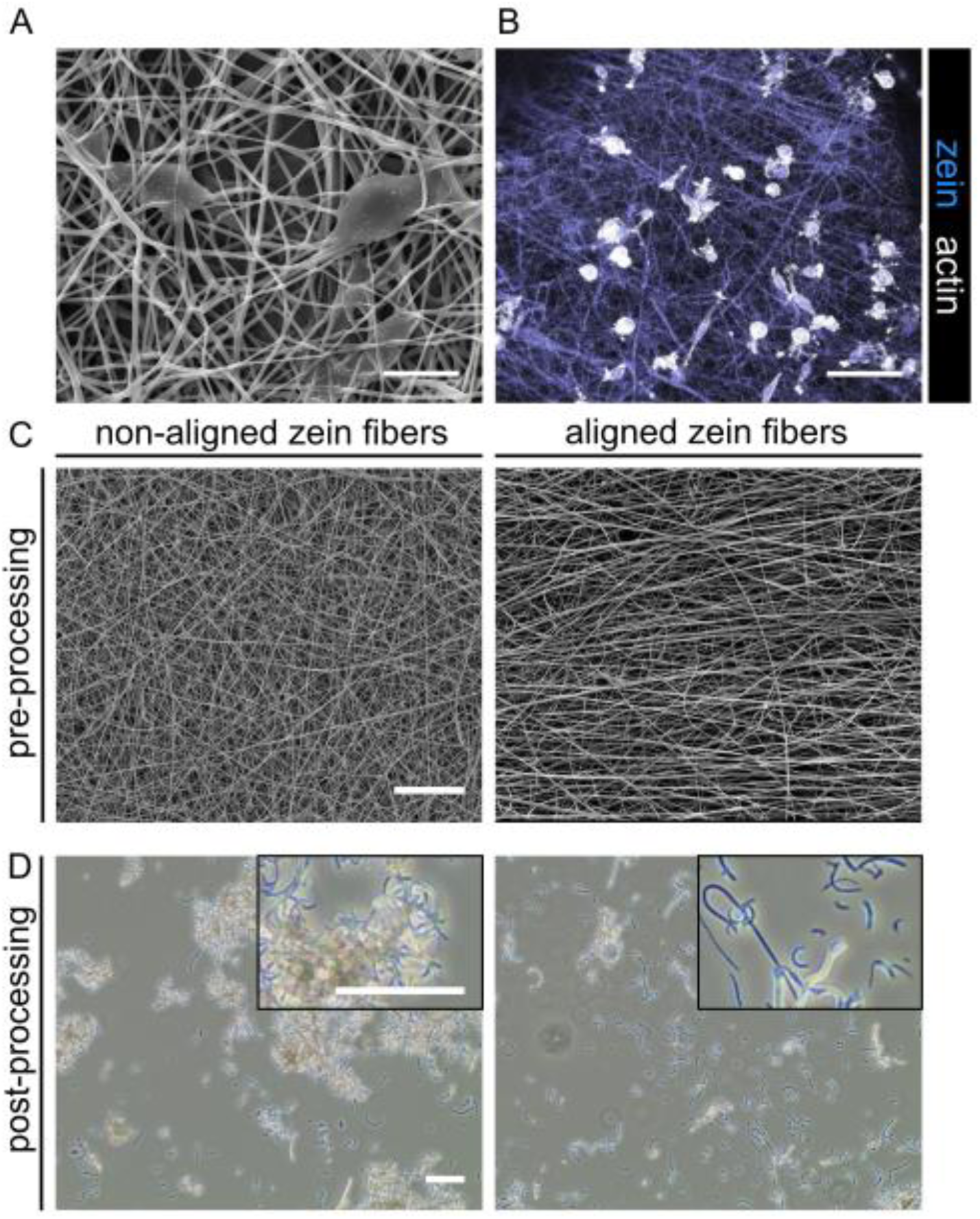
Cell attachment and processing of electrospun fibers. A: Scanning electron microscope (SEM) images showing cell attachment to zein fibers. Scale bar, 10 µm. B: Fluorescence microscopy images of satellite cells attaching to zein fibers. Phalloidin, white; Hoechst, blue. Scale bar, 100 µm. C: SEM images showing the morphology of aligned and non-aligned zein fibers before processing. Scale bar, 50 µm. D: Brightfield microscopy images of zein fibers shown in C, post-processing with ultrasonication. Scale bars, 75 µm.

### Zein fiber addition leads to faster biomaterial degradation (Fig. 4)

To investigate the effect of zein fibers on the mechanical and physical properties of an alginate hydrogel, and to understand the interactions between alginate polymers and zein fibers, we blended the SSFs at two different concentrations (0.1 and 0.5 wt%), and investigated the gel response in terms of stiffness, relative remaining mass and swelling behavior (Fig. 4a, b and c), since these properties facilitate cell migration, morphology and indirectly affect differentiation capacity.

**Figure 4:**
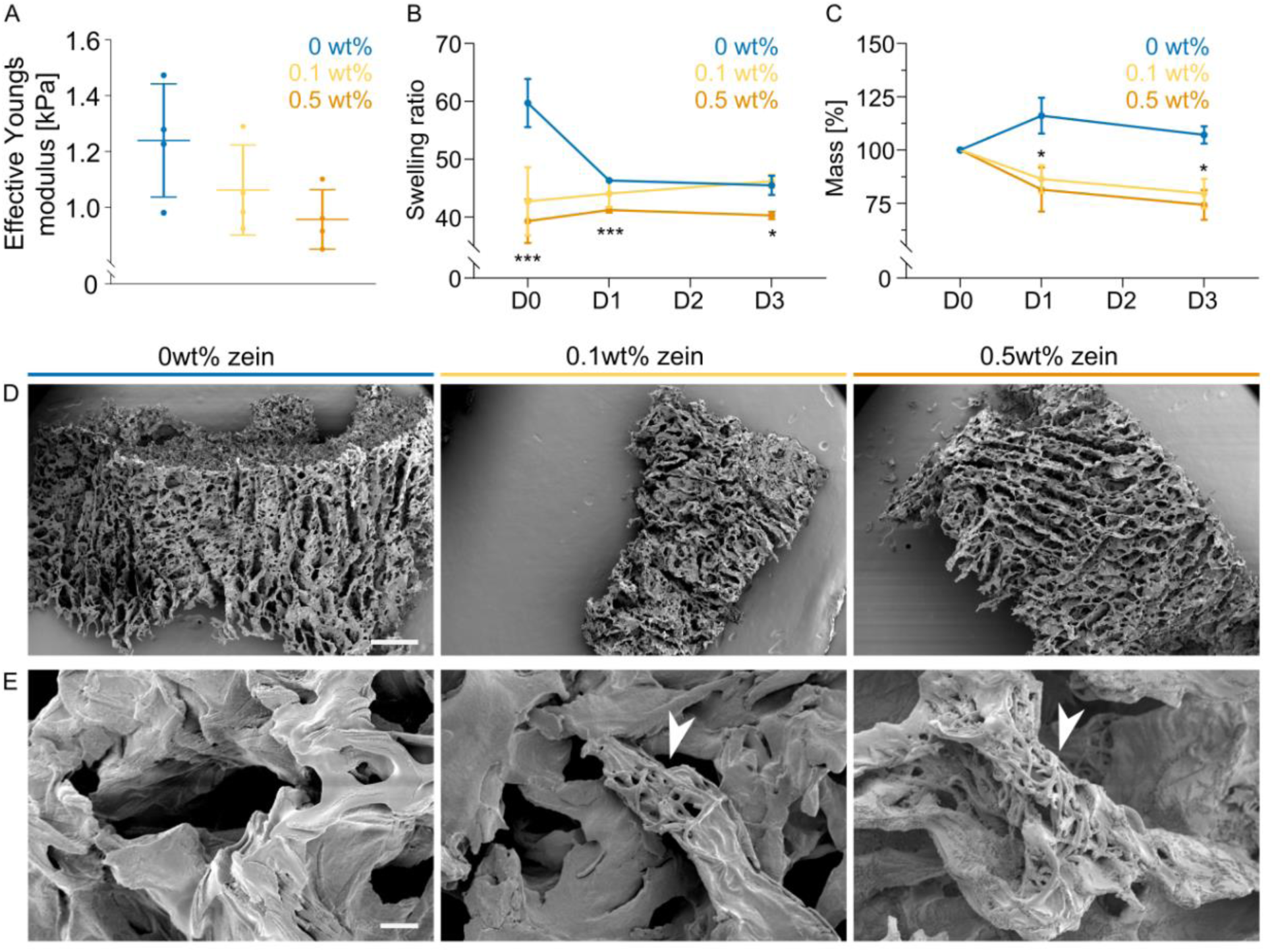
Characterization of alginate hydrogels with and without electrospun zein fibers. A: Hydrogel stiffness on day 0 with 0, 0.1 and 0.5 wt% zein-SSFs without cells.Error bars indicate s.d., n = 4. B: Swelling of hydrogels on days 0, 1 and 3 with 0, 0.1 and 0.5 wt% zein-SSFs. Error bars indicate s.d., n = 3. C: Relative remaining mass of hydrogels on days 0, 1 and 3 with 0, 0.1 and 0.5 wt% zein-SSFs. Error bars indicate s.d., n = 3. D: Scanning electron microscopy (SEM) images showing structure of alginate hydrogels with 0, 0.1 and 0.5 wt% zein-SSFs. Scale bar, 500 µm. E: SEM images showing morphology of alginate hydrogels with 0, 0.1 and 0.5 wt% short stranded zein fibers. Arrowheads indicate the presence of zein-SSFs. Scale bar, 10 µm. *P*-values; * *P* ≤ 0.05, ** *P* ≤ 0.01, *** *P* ≤ 0.001.

Alginate stiffness upon calcium crosslinking did not differ significantly, however a trend of decreasing stiffness with increasing fiber concentration was observed (Fig. 4a). Correlating with this result, we saw an enhanced degradation of the gel with increasing fiber addition (Fig. 4c). These two results in combination show that the fiber addition affects the mechanical properties which the cells sense when they get encapsulated during BAM formation. Additionally, we observed that the swelling behavior of the gel was affected by fiber addition (Fig. 4b), which could be linked to the hydrophobicity of zein.

### Addition of zein fibers improves compaction during cultured muscle formation (Fig. 5)

After addition of zein fibers to the alginate, FHCs were tested in a 3D tissue culture system. The first step for tissue formation in a self-assembling hydrogel is cell-driven compaction^22^. The majority of observed compaction occurred between days 0 and 3, where we also observed the most significant differences between the three groups (Figs. 5a, b). While the final extent of compaction was not affected by the addition of zein fibers, the rate of BAMs compaction was substantially increased. On day 1, BAMs incorporating 0.1 and 0.5 wt% zein-SSFs showed significantly increased compaction when compared to the control group (Figs. 5a, b).

**Figure 5:**
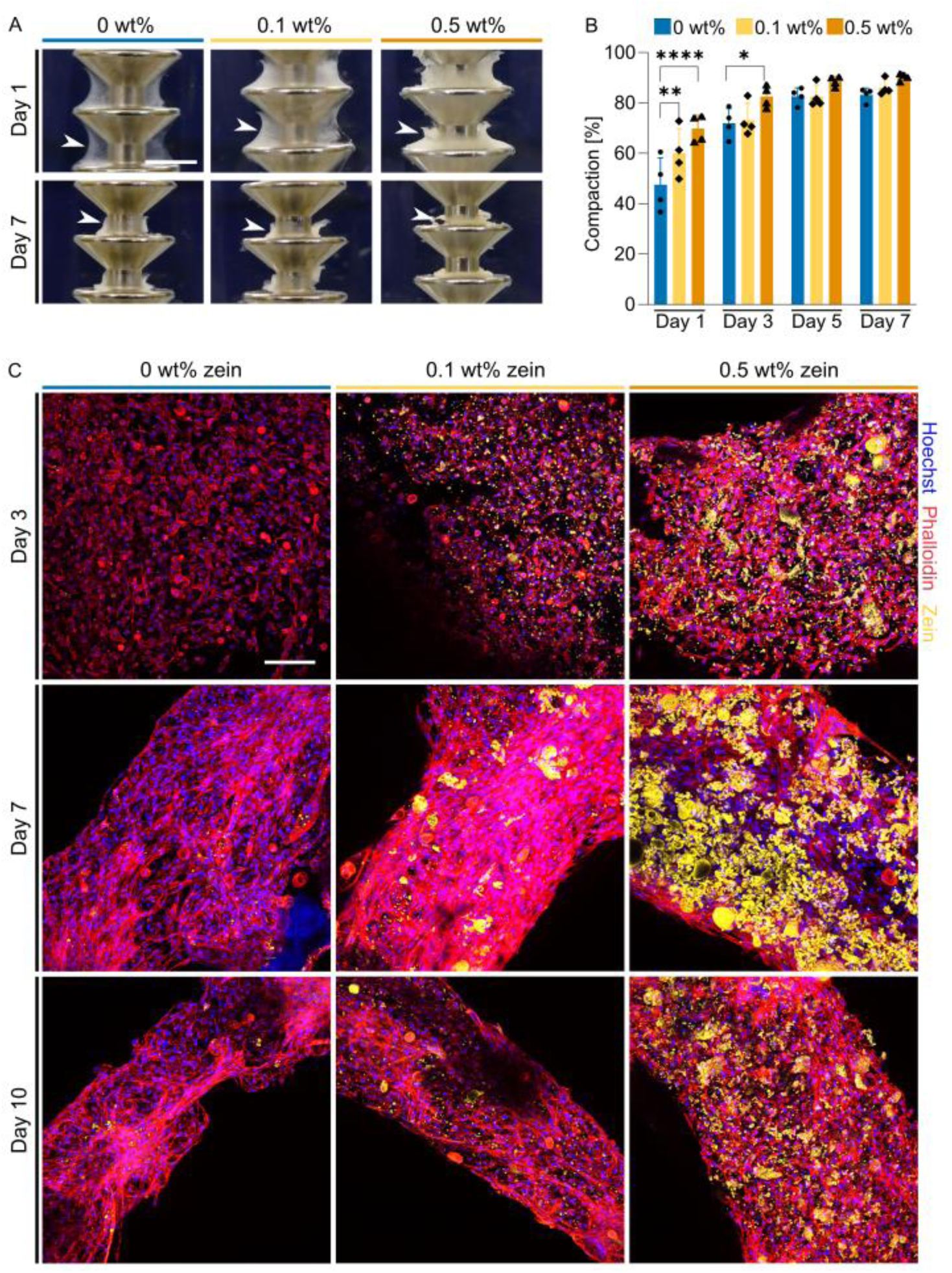
Blending of short-stranded zein fibers improves compaction during cultured muscle formation. A: BAM formation around stainless steel pillars after 1 and 7 days culture with 0, 0.1 and 0.5 wt% short stranded zein fibers. Arrowhead’s indicate the position of the BAMs around the pillar. Scale bar, 3 mm. B: Quantification of images shown in (a) with additional timepoints. Degree of compaction on days 1, 3, 5 and 7. Error bars indicate s.d., n = 4. C: Fluorescent images showing BAM morphology on days 3, 7 and 10. Phalloidin, red; zein, green; Hoechst, blue. Scale bar, 100 µm*. P*-values; * *P* ≤ 0.05, *P* ≤ 0.01, *** *P* ≤ 0.001.

To investigate the structure and distribution of zein-SSFs within the BAM construct, and the interaction of cells with the incorporated SSFs in more detail, we coupled the zein with a green fluorescent dye. After fixing 3D muscle tissues on days 3, 7 and 10, we could visualize spreading and alignment of the cells via F-actin staining, which was enhanced with the addition of 0.1 wt% zein (Fig. 5c). Addition of 0.5 wt% zein-SSFs, on the other hand, showed decreased F-actin expression and reduced cell alignment. The incorporated SSFs did not maintain their fibrous structure throughout the culture period, instead starting to cluster into spherical structures, which subsequently degraded over time (Fig. 5c).

### Zein fiber addition improves protein yield and metabolic activity (Fig. 6)

Having examined morphological changes during the course of BAM cultures, we next looked into cellular metabolic activity, including accumulation of protein in order to assess the extent of myogenic differentiation. Total protein measurements showed protein accumulation in both control and 0.1 wt% zein, while the 0.5 wt% zein group showed the highest total protein yield on day 3, which didn’t accumulate further on days 7 and 10. The 0.1 wt% zein group showed increased protein accumulation on days 7 and day 10 compared to the control group (Fig. 6a).

**Figure 6:**
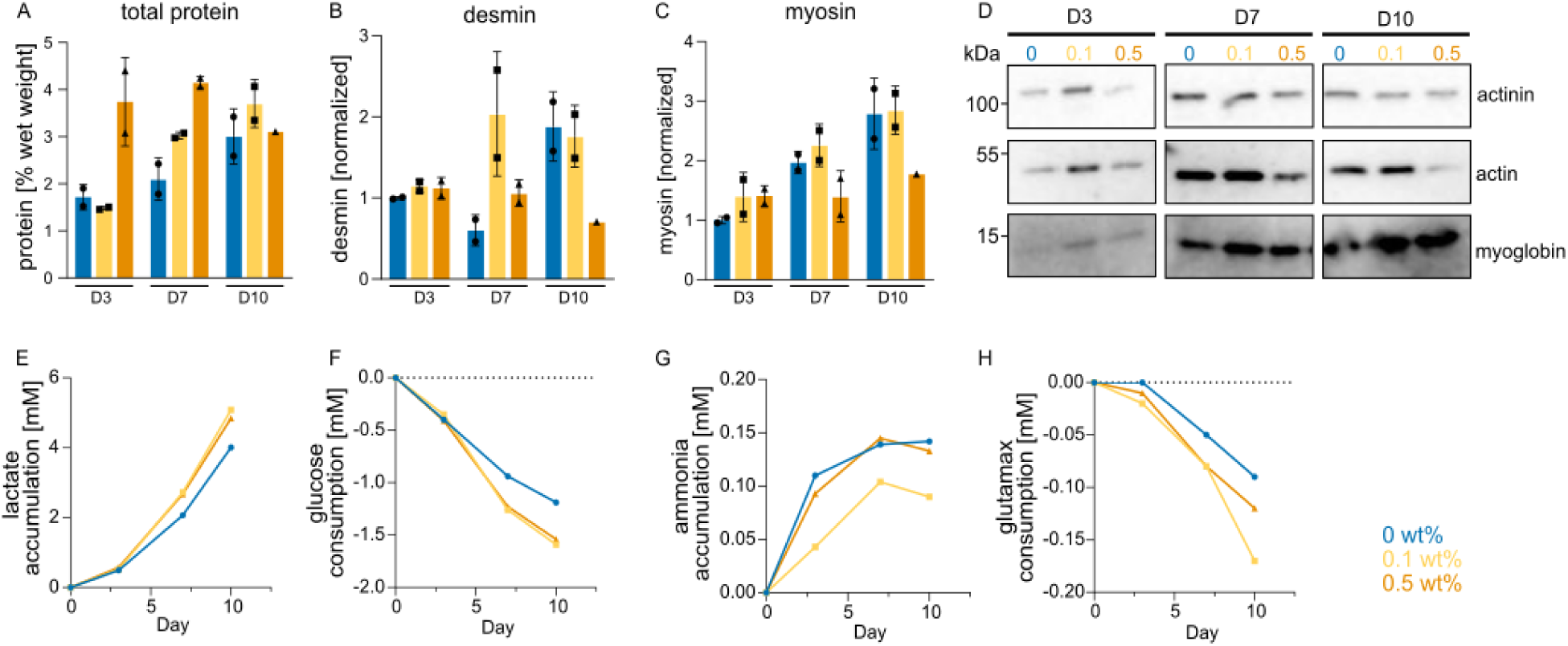
Protein yield and metabolic changes after zein fiber addition. A: Total protein measurements for BAMs with 0, 0.1 and 0.5 wt% zein fibers at indicated time points. Error bars indicate s.d., n = 2. No statistically significant differences were observed, with *P*-values > 0.05. B: Desmin ELISAs in BAMs with 0, 0.1 and 0.5 wt% zein fibers. Error bars indicate s.d., n = 2. *P*-values > 0.05. C: As B, but slow myosin heavy chain ELISAs. Error bars indicate s.d., n = 2. *P*-values > 0.05. D: Western blot of selected muscle-related proteins during myogenic differentiation, analyzed at indicated time points. E: Lactate concentrations recorded in culture medium at indicated time points. Values are normalized to day 0. *P*-values > 0.05. F: As E, but for glucose concentrations. G: As E, but for ammonia concentrations. H: As E, but for glutamax concentrations.

To quantitatively compare the changes in muscle differentiation through the addition of zein fibers, a selection of muscle specific proteins was measured. Desmin was chosen as a marker for early fusion while myosin heavy chain showed the progression in the differentiation (often referred to as maturation). The group with 0.1 wt% zein-SSFs showed an earlier increase in both desmin and myosin heavy chain compared with the control group (Figs. 6b, c). The group with 0.5 wt% zein fibers showed significantly reduced expression of muscle proteins, which also correlated with the reduced actin fibers and alignment previously observed (Figs. 4c, 6b, 6c). Other muscle-specific proteins such as alpha-actin-1, alpha-actinin, and myoglobin were measured by Western blot analysis and showed similar trends to the ELISAs (Fig. 6d).

To further examine metabolic activity of the myoblasts, glucose and glutamax consumption were measured, as well as the respective accumulation of lactate and ammonia (Figs. 6e-h). Overall the metabolic activity of the cells was increased by zein-SSFs addition, suggesting that the construct viability was increased leading to longer-lasting tissues (Figs. 6e-h). Contrary to the glucose, glutamax and lactate results, ammonia accumulation was reduced in the 0.1 wt% zein fiber group (Fig. 6g).

## Discussion

The primary objective of this study was to facilitate assembly of skeletal muscle tissue in an alginate hydrogel system by improving compaction and structural stability as a prerequisite for enhanced myogenic differentiation and maturation. Our approach involved the development of fiber hydrogel composites (FHCs), through incorporation of short stranded zein fibers into alginate hydrogels, with the aim of mimicking the stiffness and structural features of native ECM and extensively studied animal-derived hydrogels.

As a first step to making alginate FHCs, we fabricated and fragmented electrospun fibrous zein scaffolds, based on previously described methods^21,23,24^. Successful formation of a stable Taylor cone (also known as ‘electrohydrodynamic jet’) was contingent upon optimization of parameters such as viscosity, conductivity and surface tension of the spinning solution, in combination with the applied voltage^25^. Electrospinning of zein, due to its hydrophobic properties, commonly employs acetic acid and organic solvents, and the choice of solvent is known to influence the electrospinning process through the different degree of protein unfolding they induce^26,27^. To illustrate this, Selling *et al*. have shown that zein fibers electrospun from alcoholic solutions adopt a ribbon-like morphology, while fibers electrospun with acetic acid solutions resulted in more rounded fiber morphologies at similar concentrations^27^. This can be explained by the lower vapor pressure of acetic acid, which reduces the evaporation at the needle tip and thus allows for more robust round fiber formation^27,28^. In the present study, we optimized the spinning with a mixture of both acetic acid and ethanol. Spinning solutions with 20 and 25 wt% zein failed to form Taylor cones, and thus led to heterogenous fiber formation, due to their low viscosity and consequently low degree of polymer entanglement^27^. While increased viscosity and reduced conductivity stabilized Taylor cone formation, 30 and 35 wt% zein solutions caused needle clogging. However, homogenous fibers of approximately 600 nm diameter could be achieved with 28 wt% zein solution.

Following ultrasonication, zein fibers were homogeneously incorporated into alginate hydrogels by blending two different final concentrations (0.1 and 0.5 wt%). After zein fiber addition, we observed disparities in swelling and degradation behaviors. Immediately after gel formation, swelling was significantly reduced, which can be interpreted as a function that is introduced by the hydrophobicity of the zein fibers. Mao *et al*. similarly observed that mixing hydrophobic and fragmented PCL fibers into alginate/gelatin hydrogels resulted in a reduced swelling behavior compared to hydrogels alone^19^. While showing trends towards reduced stiffness with increasing fiber concentration, no significant differences could be observed between groups. Degradation was enhanced upon increasing zein fiber concentration, suggesting that the secondary component addition interfered with the intermolecular interactions between alginate polymer chains involving divalent calcium ions (Fig. S1). Manufacturing electrospun fibers from alginate zein mixtures would circumvent this, by improving the interaction between SSFs and hydrogel network.

The enhanced degradation concurs with differences we observed in early tissue assembly, timing and extent of compaction behavior, and actin fiber formation. Compaction of BAMs usually occurs within the first 72 h after cell encapsulation, and is based on both alginate degradation dynamics and cell-driven tissue reorganization as cells interact with the biomaterial blend. Within the initial 72 h timeframe, BAMs displayed an enhanced compaction rate and increased degradation due to zein-SSF presence. Consequent increase in tension, similar to collagen-based systems, lead to improved actin fiber assembly and alignment. Notably, cell alignment and tension have previously been linked to enhanced myogenic differentiation^29–31^. Yeo *et al*. demonstrated this by showing higher degree of myotube alignment and myogenic differentiation on aligned PCL/alginate scaffolds compared to non-aligned scaffolds^30^. Similarly, we observed that the presence of 0.1 wt% zein-SSF improved actin protein expression, accompanied by tension build-up around the stainless-steel pillar, leading to improved early-stage differentiation.

It is desirable for the main protein fraction of a cultivated meat product to be cell-derived in order to achieve meat mimicry, but, ultimately, a balance must be struck between structural stability for cell guidance and a minimum quantity of plant-based material in the final product. The improvements in tissue formation and alignment lead to more stable and durable constructs needed for maturation and thus protein accumulation, which is important for the quality and cost efficiency of the final product. As previously mentioned, texture and nutritional value are achieved by sufficient maturation of the muscle fibers. Furthermore, taste, cooking behavior and water retention are key factors that determine cultivated meat quality. For the cooking behavior and final taste, the Maillard reaction plays a key role, and can be further enhanced by resident zein fibers. Furthermore, the hydrophobicity of zein could play an important role in the water holding capacity of the cultivated meat product.

Overall, the incorporation of zein-SSFs into alginate-based hydrogels is a promising system to improve muscle tissue formation and increase BAM construct longevity.

## Material & Methods

### Preparation and characterization of spinning solutions and fibers

#### Solution preparation

Zein (Flozein; #F4400) was dissolved in concentrations of 20, 25, 28, 30 and 35 wt% in a ratio of 1:1 (v/v) ethanol:acetic acid. Solutions were stirred for 1 h at room temperature prior to electrospinning and fluorescent zein was added in 1:100 dilution.

#### Viscosity measurements

Dynamic viscosity (mPa.s) of zein solutions at 25 °C was determined with a rotational viscometer (ViscoQC 300, Anton Paar GmbH) equipped with a concentric cylinder measuring system (CC26). Measuring volume was 10 mL with a minimum torque of 10%. A constant rotational speed was used to determine the dynamic viscosity value with 20, 10 and 5 rpm were used for the 20-28, 30 and 35% zein solutions respectively.

#### Conductivity measurements

Conductivity (μS cm^-1^) of zein solutions at room temperature was determined with a SevenCompact pH/Ion meter (S220, Mettler Toledo) equipped with a conductivity probe.

#### Electrospinning

Zein solutions were charged to 25 kV with a high voltage power supply (IME). Solutions were spun at a flow rate of 2 ml h^-1^ and fibers were collected on a grounded rotating stainless steel cylinder with a distance of 90 mm between needle and collector. Experiments were carried out at 25 °C and 40% relative humidity.

#### Scanning electron microscopy

Fiber morphology was analyzed using a scanning electron microscope (JSM-6010LV Plus, Jeol). Fibers were sputtered with gold prior to imaging (JFC-1300 10BA, Jeol). Average fiber diameters were measured using ImageJ, taking the average of 100 fibers for each sample.

#### Ultrasonication

Electrospun meshes were immersed in PBS at a concentration of 10 mg ml^-1^. Samples were treated at 4 °C for 30 min at 37 kHz (Elmasonic P 120 H, Elma).

#### Fluorescent-labeled zein

Zein was fluorescently labeled by reacting its lysine residues with the fluorescent dye derivative Atto 488 NHS ester (ATTO-TEC, AD 488-31). Zein was dissolved in ethanol at 10 wt% and 50 mM Atto 488 NHS ester was added and stirred overnight in the dark.

### Hydrogel formation and 3D cell culture

#### Cell isolation and purification

Cells were isolated from semitendinosus muscles of Belgian Blue cattle by collagenase digestion (CLSAFA, Worthington; 1 h; 37 °C), filtration with 100 μm cell strainers, red blood lysis using a Ammonium-Chloride-Potassium (ACK) lysis buffer, and final filtration with 40 μm cell strainers. Cells were plated in serum-free growth medium (SFGM, Supplementary Table 1) on fibronectin-coated (4 μg cm^-2^ bovine fibronectin; F1141, Sigma-Aldrich) tissue culture vessels. SCs were purified at 72h post-isolation using MACSQuant Tyto Cell Sorter (Miltenyi Biotec), using a cell surface marker panel of ITGA5 and ITGA7 (Supplementary Table 2), as previously described^32^.

#### Synthesis of RGD-functionalized alginate

Sodium alginate (L3; KIMICA) was purified via dialysis (3.5-kDa membrane) at a 1 wt% concentration in water for 3 days at room temperature. The water was replaced daily and the solution was lyophilized (L-200, Buchi) to obtain the solid material. Next, purified alginate was dissolved in 100 mM MES and 300 mM NaCl buffer at pH 6.5 to create a 1 wt% solution. Once dissolved, N-hydroxysulfosuccinimide sodium salt (sulfo-NHS; TCI) and 1-Ethyl-3-(3-dimethylaminopropyl) carbodiimide hydrochloride (EDC-HCl; Sigma-aldrich) were added at a concentration of 0.47 and 0.95 mg mL^-1^, respectively. After 30 min, pH was adjusted to 8.0, NH_2_-G4RGDSP-COOH (Biomatik) was added at a concentration of 1 μg mL^-1^ and allowed to react for 20 h. The reaction mixture was purified in 100, 50, 25 and then 0 mM NaCl for 3 days, replaced daily and finally lyophilized to obtain the RGD-functionalized alginate product.

#### Gel degradation assays

Purified and lyophilized alginate prepared at a concentration of 0.9 wt% in PBS (Gibco; #10010023) was mixed with short stranded zein fibers at concentrations of 0.1 wt% and 0.5 wt% and crosslinked in 100 mM CaCl_2_ for 5 min. Crosslinked hydrogels were washed once and incubated in PBS at 37 °C for a maximum of 72 h. Samples were taken at 0, 24 and 72 h. After PBS removal, wet weight was measured and post-lyophilization, dry mass was determined at the respective timepoints.

#### Mechanical properties

Effective Young’s moduli were determined using a PIUMA nanoindenter (Optics11, Amsterdam, Netherlands) on crosslinked hydrogels. Hydrogels were crosslinked around stainless steel pillars as shown in Fig. 4. Gels were maintained in a buffer containing 50 mM CaCl_2_, 50 mM 3-(N-morpholino)propanesulfonic acid and 150 mM NaCl. Indentations were performed using a probe with a tip radius of 51 μm and cantilever stiffness of 0.43 N m^-1^. Effective Young’s modulus was determined by fitting a Hertzian Contact model over 80% of the load-indentation curve.

#### 3D cell culture

RGD-functionalized alginate was dissolved at a concentration of 1.8 wt% in PBS and mixed with short stranded zein fibers (0, 0.1 & 0.5 wt%). Bovine SCs were expanded in SFGM (Supplementary Table 1) and after harvesting resuspended at 1.2 x 10^7^ cells ml^-1^ in serum-free differentiation medium (SFDM Supplementary Table 1). Cell suspension and alginate/zein solutions were mixed at a 1:1 ratio. Gel mixtures were injected around stainless steel pillars and crosslinked by immersion in 100 mM CaCl_2_ for 5 min. After crosslinking, BAMs were washed and incubated in SFDM. Medium was changed on day 3 and 7, and from day 3 onwards, SFDM was additionally supplemented with 10 µM acetylcholine. After a total of 10 days, BAMs were harvested for protein quantification and confocal microscopy. For compaction assays, BAMs were imaged every day during the 7 day period (DC-TZ90, Panasonic Lumix). Images were analyzed in ImageJ.

#### Protein measurements

Protein was extracted using RIPA Lysis Buffer (sc-24948, Santa Cruz Biotechnology) after freezing at −20 °C and lyophilization. Total protein was measured using BCA assay (ThermoFisher, 23235).

#### ELISAs

Protein extracts were tested for relative desmin and slow myosin heavy chain (MYH7) content using in-house ELISAs. For desmin ELISA, samples were diluted to 20 µg ml^-1^ total protein in carbonate coating buffer (pH 9.5) supplemented with 5% RIPA and 100 µg ml^-1^ bovine serum albumin (BSA), and added a Nunc Maxisorp plate (Invitrogen 44-2404-21). After a 2 h incubation period, wells were washed and blocked with 10 mg ml^-1^ BSA in TBS. Samples were stained with indicated primary and respective secondary antibodies (Supplementary Table 2). For myosin ELISAs, plates were incubated overnight with a capture antibody (Supplementary Table 2) in TBS at 4 °C. After blocking, samples diluted to 100 µg ml^-1^ protein in TBS containing 0.1% Tween-20 and 100 µg ml^-1^ BSA were added to the pre-coated and pre-blocked ELISA plates. Following 2 h incubation, biotinylated detection antibody was added (biotinylation of carrier and preservation-free antibody was performed with the Abcam Lightning Link biotinylation kit, ab201795), respective secondary antibody was added (Supplementary Table 2). After final wash, signals of both ELISAs were visualized using 1-step Ultra TMB ELISA substrate (Fisher Scientific 34028), an incubation period at RT and addition of 2 M sulfuric acid as the stop reagent. Absorbance values at 450 nm were measured using a plate reader (Victor X-V, Perkin Elmer).

#### Western blotting

Protein samples were boiled in reducing Laemmli buffer (5 min), separated by SDS-PAGE and transferred to PVDF membranes by electrophoresis. Equal protein loading was confirmed by full-protein staining using Revert 700 Total protein stain (LI-COR; 926-11015). Blots were stained with primary antibodies against indicated muscle specific proteins and respective secondary antibodies (Supplementary Table 2). Blots were developed using SuperSignal West Femto Maximum Sensitivity Substrate (Thermo Scientific) and visualized on an Azure 600 chemiluminescence imager (Azure Biosystems).

#### Tissue staining

BAMs were fixed with 4% paraformaldehyde (PFA) in a wash buffer containing 154 mM NaCl, 50 mM CaCl_2_ and 50 mM 3-(N-Morpholino)propanesulfonic acid, permeabilized with 0.5% Triton X-100, and blocked in 5% bovine serum albumin (BSA). Cells were stained for F-actin (Supplementary Table 2) and Hoechst 33342 (Thermo Scientific). Samples were imaged using a confocal microscope (TCS SP8, Leica Microsystems)

#### Statistical analysis

Statistical significance was assessed using Prism v9.3.1 (GraphPad). Two-way analysis of variance (Figs. 3b-c, 4b, 6a-c and 6e-h) was performed to determine statistical significance. Adjusted *P*-values used throughout the figures were: * *P* ≤ 0.05, ** *P* ≤ 0.01, *** *P* ≤ 0.001.

## Data availability

Data supporting the findings of this study are available from the authors on request.

## Acknowledgements

We would like to thank Paul Wieringa and Shivesh Anand for providing help during the electrospinning optimization process. Additionally, we would like to thank Medace B.V. for use of their lab spaces and facilities.

## Author contributions

LM, SS, NH and AJGP performed experiments and analysis. SS, JEF, MJP and AD supervised the study. LM and AD wrote the manuscript with input from all authors.

