## Supplementary Material for "Short-stranded zein fibers for muscle tissue engineering in alginate-based hydrogels"

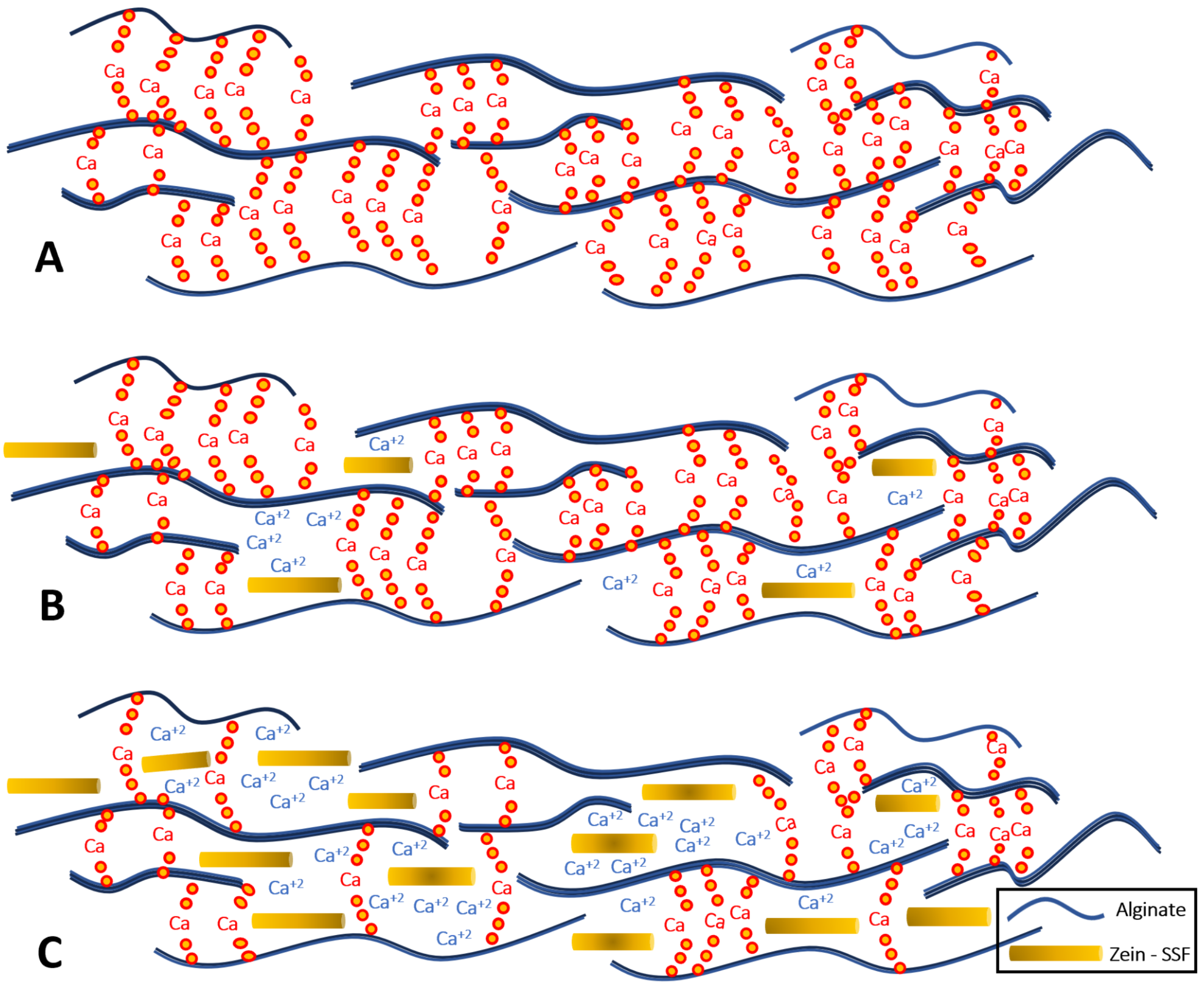


**Supplementary Figure 1: Zein interference of calcium crosslinking in alginate hydrogels.**

A: Without zein-SSF addition, uninterrupted ionic crosslinking

B: low concentration of zein-SSFs, with decreased ionic crosslinking

C: high concentration of zein-SSFs, with further decreased ionic crosslinking

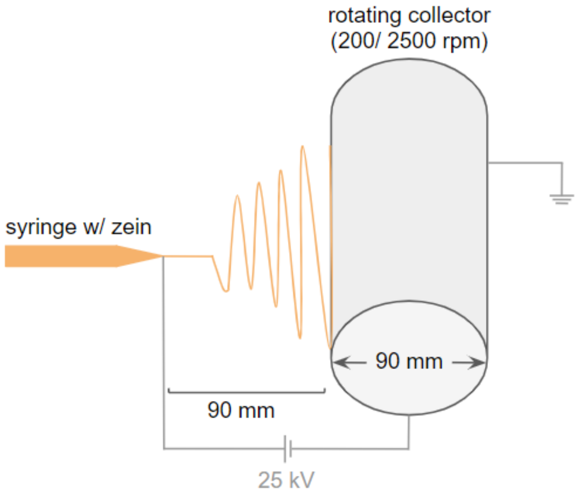


**Supplementary Figure 2: Electrospinning set up**

**Supplementary Table 1: Media formulations.**

| # | Component | Reference | Concentration |
|---|---|---|---|
| **Serum-free growth medium (SFGM)** | | | |
| 1 | DMEM/F-12 | P04-041262B, PAN Biotech | |
| 2 | α-linolenic acid | L2376, Sigma Aldrich | 1 μg $ml^{-1}$ |
| 3 | bFGF-2 | 100-18B, Peprotech | 10 ng $ml^{-1}$ |
| 4 | bHGF | 100-39H, Peprotech | 50 ng $ml^{-1}$ |
| 5 | Bovine Serum Albumin (BSA) | A9418, Sigma Aldrich | 5 mg $ml^{-1}$ |
| 6 | D-glucose | G7021, Sigma Aldrich | 17.7 mM |
| 7 | Glutamax | 35050061, ThermoFisher | 2 mM |
| 8 | Hydrocortisone | H0888, Sigma Aldrich | 36 ng $ml^{-1}$ |
| 9 | IGF-1 | 100-11, Peprotech | 100 ng $ml^{-1}$ |
| 10 | ITSE | 00-101, biogems | 1% |
| 11 | L-ascorbic acid 2-phosphate (Vitamin C) | A8960, Sigma Aldrich | 155 μM |
| 12 | LIF | | 5 ng $ml^{-1}$ |
| 13 | PDGF-BB | 100-14B, Peprotech | 10 ng $ml^{-1}$ |
| 14 | Penicillin/Streptomycin/Amphotericin (PSA) | 17-745E, Lonza | 1% |
| 16 | VEGF | 100-20, Peprotech | 10 ng $ml^{-1}$ |
| **Serum-free myogenic differentiation medium (SFDM)** | | | |
| 1 | DMEM | A14430-01, Gibco | |
| 2 | EGF-1 | AF-100-15, Peprotech | 10 ng $ml^{-1}$ |
| 3 | D-glucose | G7021, Sigma | 5.5 mM |
| 4 | GlutaMax | 35050061, ThermoFisher | 2 mM |
| 5 | Human Serum Albumin | Rc HA NW20, Richcore Lifesciences | 0.5 mg $ml^{-1}$ |
| 6 | ITSE | 00-101, biogems | 2% |
| 7 | L-ascorbic acid 2-phosphate (Vitamin C) | A8960, Sigma Aldrich | 40 μM |
| 8 | MEM Amino Acids Solution | 11130-051, ThermoFisher | 0.50% |
| 9 | $NaHCO_3$ | P2256, Sigma Aldrich | 6.5 mM |
| 10 | Penicillin/Streptomycin/Amphotericin (PSA) | 17-745E, Lonza | 1% |
| 11 | Soy hydrolysates | 58903C, Merck | 1% |
| 12 | Sodium l-lactate | 71718, Sigma | 10 mM |
| 13 | Sodium pyruvate | P2256, Sigma Aldrich | 0.5 mM |

**Supplementary Table 2: Antibodies used in this study.**

| Target | Colour | Source | Dilution | Reference | Application |
|---|---|---|---|---|---|
| α-alpha-actin-1 | - | Abcam | 1:5000 | ab184705 | Western Blot |
| α-actinin | - | Sigma-Aldrich | 1:2500 | A7811 | Western Blot |
| desmin | - | Abcam | 1:3000 | ab227651 | ELISA |
| f-actin | Atto550 | Sigma-Aldrich | 1:300 | 19083 | IF |
| ITGA5 | PE | Miltenyi Biotec | 1:50 | 130-110-532 | Flow |
| ITGA7 | APC | Miltenyi Biotec | 1:50 | 130-123-833 | Flow |
| myoglobin | - | Abcam | 1:2500 | ab231725 | Western Blot |
| myosin | - | Abcam | 1.5 $\mu g\ ml^{-1}$ | ab11083 | ELISA |
| myosin | - | Abcam | 0.8 $\mu g\ ml^{-1}$ | ab197687 | ELISA |
| streptavidin | HRP | Abcam | 1:1000 | ab7403 | ELISA |
| anti-mouse | HRP | Dako | 1:2000 | P0447 | Western Blot |
| anti-rabbit | HRP | Abcam | 1:8000 | ab6721 | ELISA, Western Blot |
